# Decomposing information into copying versus transformation

**DOI:** 10.1101/584771

**Authors:** Artemy Kolchinsky, Bernat Corominas-Murtra

**Affiliations:** Santa Fe Institute, 1399 Hyde Park Road, Santa Fe, NM 87501, USA; Institute of Science and Technology Austria, Am Campus 1, A-3400, Klosterneuburg, Austria

## Abstract

In many real-world systems, information can be transmitted in two qualitatively different ways: by *copying* or by *transformation. Copying* occurs when messages are transmitted without modification, e.g., when an offspring receives an unaltered copy of a gene from its parent. *Transformation* occurs when messages are modified systematically during transmission, e.g., when non-random mutations occur during biological reproduction. Standard information-theoretic measures do not distinguish these two modes of information transfer, although they may reflect different mechanisms and have different functional consequences. Starting from a few simple axioms, we derive a decomposition of mutual information into the information transmitted by copying and by transformation. Our decomposition applies whenever the source and destination of the channel have the same set of outcomes, so that a notion of message identity exists, although generalizations to other kinds of channels and similarity notions are explored. Furthermore, copy information can be interpreted as the minimal work needed by a physical copying process, relevant to better understand the physics of replication. We use the proposed decomposition to explore a model of amino acid substitution rates. Our results apply to any system in which the fidelity of copying, rather than simple predictability, is of critical relevance.

## I. INTRODUCTION

Shannon’s information theory provides a powerful set of tools for quantifying and analyzing information transmission. A particular measure of interest is *mutual information*, which is the most common way of quantifying the amount of information transmitted from a source to a destination. Mutual information has fundamental interpretations and operationalizations in a variety of domains, ranging from telecommunications [1, 2], gambling and investment [3–5], biological evolution [6], statistical physics [7, 8], and many others. Nonetheless, it has long been observed [9, 10] that mutual information does not distinguish between a situation in which the destination receives a *copy* of the source message versus one in which the destination receives some systematically *transformed* version of the source message (where by “systematic” we refer to transformations that do not arise purely from noise).

As an example of where this distinction matters, consider the transmission of genetic information during biological reproduction. When this process is modeled as a communication channel from parent to offspring, the amount of transmitted genetic information is often quantified by mutual information [11–15]. However, during reproduction, genetic information is not only copied but can also undergo systematic transformations in the form of “nonrandom mutations” [16, 17]. While mutual information does not distinguish which part of genetic information is transmitted by exact copying and which part is transmitted by nonrandom mutations, these two modes of information transmission are driven by different mechanisms and have dramatically different evolutionary and functional implications, not least because nonrandom mutations typically lead to deleterious or even fatal consequences.

The goal of this paper is to find a general decomposition of the information transmitted by a channel into contributions from copying versus from transformation. In Fig. 1, we provide a schematic that visually illustrates the problem. Essentially, we seek a decomposition of transmitted information into copy and transformation that distinguishes the example provided in (Fig. 1a), where the copy is perfect, from the one provided in (Fig. 1b), where the message has been systematically scrambled, from the one provided in (Fig. 1c), where the channel is completely noisy. Of course, we want also such a decomposition to apply in less extreme situations, where part of the information is copied and part is transformed.

**Figure 1.**
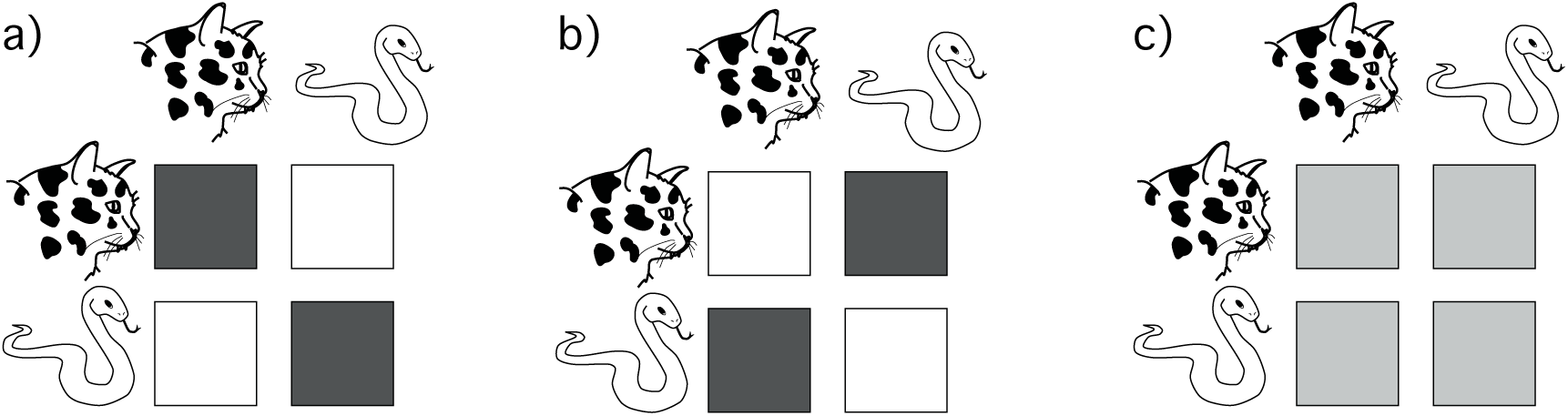
An illustration of the problem of copy and transformation. Consider three channels, each of which can transmit two messages, indicated by *cat* and *snake* –e.g., alarm calls in an animal communication system. In all panels, the rows indicate the message selected at the source, the columns indicate the message received at the destination, and the shade of the respective square indicates the conditional probability of the destination message given source message. For the channel in (a), all information is copied: the channel maps *cat* → *cat* and *snake* → *snake* with probability 1. For the channel in (b), all information is transformed: the channel maps *cat* → *snake* and *snake* → *cat* with probability 1. Note that for any source distribution, the mutual information between source and destination is the same in (a) and (b). The channel in (c) is completely noisy: the probability of receiving different messages at the destination do not depend on the message selected at the source, and the mutual information between source and destination is 0. Observe that transformation is different than noise, in it still involves the transmission of information.

The distinction between copying and transformation is important in many other domains beyond the case of biological reproduction outlined above. For example, in many models of animal communication and language evolution, agents exchange signals across noisy channels, and then use these signals to try to agree on common referents in the external world [10, 18–24]. In such models, successful communication occurs when information is transmitted by copying; if signals are systematically transformed — e.g., by scrambling — the agents will not be mutually intelligible, even though mutual information between them may be high. As another example, the distinction is relevant in the study of information flow during biological development, where recent work has investigated the ability of regulatory networks to decode development signals, such as positional information, from gene expression patterns [25]. In this scenario, information is copied when developmental signals are decoded correctly, and transformed when they are systematically decoded in an incorrect manner. Other examples are provided by Markov chain models, which are commonly used to study computation and other dynamical processes in physics [26], biology [27] or sociology [28], among other fields. Indeed, Markov chains can be seen as a communication channel in which the system state transmits information from the past into the future. In this context, copying occurs when the system maintains its state constant over time (remains in fixed points) and transformation occurs when the state undergoes systematic changes (e.g., performs some kind of non-trivial computations).

Interestingly, while the distinction between copy and transformation information seems natural, it has not been previously considered in the information-theoretic literature. This may be partly due to the different roles that information theory has historically played: on one hand, a field of applied mathematics concerned with the engineering problem of optimizing information transmission — its original purpose — on the other a set of quantitative tools for analyzing intrinsic properties of real-world systems. Because of its origins in engineering, much of information theory, including Shannon’s channel-coding theorem, which established mutual information as a fundamental measure of transmitted information [2, 29, 30], is formulated under the assumption of an external agent who can appropriately encode and decode information for a given communication channel, in this way accounting for any transformations performed by the channel. However, in many real-world systems, there is no additional external agent who codes for the channel [10, 31], and one is interested in the ability to copy information across a channel without any additional encoding or decoding. This latter property is the main subject of this paper.

A final word is required to motivate our information-theoretic approach. It is standard to characterize the ability of a channel to copy messages via the “probability of error” statistic, which we indicate as *ϵ* [2]. In particular, *ϵ* reflects the probability that the destination receives a different message than the one that was sent by the source, while 1 −*ϵ* reflects the probability that the destination receives the same message as was sent by the source. However, for our purposes, this approach is insufficient. First of all, while 1 −*ϵ* quantifies the propensity of a channel to copy information, *ϵ* does not measure propensity to transmit information by transformation, since it increases both in the presence of transformation and in the presence of noise (in other words, *ϵ* is high both in channels like Fig. 1b and channels Fig. 1c). Among other things, this means that 1 −*ϵ* and *ϵ* cannot be used to compute a channel’s “copying efficiency” (i.e., which portion of the total information transmitted across a channel is copied). Second, and more fundamentally, *ϵ* and 1 −*ϵ* are not information-theoretic quantities, in the sense that they do not reflect the *amount* of information. For instance, 1 −*ϵ* is bounded between 0 and 1 for all channels, whether considering a simple binary channel or a high-speed fiber-optic line — in the language of physics, *ϵ* is an intensive property, rather than extensive one that scales with the size of the channel. We instead seek measures which quantify the amount of copied and transformed information, and which can grow as the capacity of the channel under consideration increases.

We present a decomposition of information that distinguishes copied from transformed information. We derive our decomposition by proposing four natural axioms that copy and transformation information should satisfy, and then identifying the unique measure that satisfy these axioms. Our resulting measure is easy to compute and can be used to decompose either the total mutual information flowing across a channel, or the specific mutual information corresponding to a given source message, or an even more general measure of acquired information called *Bayesian surprise*.

The paper is laid out as follows. We present our approach in the next section. In Section III, we demonstrate that copy information is related to, though different from, the notion of *rate-distortion* in information theory [2]. We also show that while our basic decomposition is defined for discrete-state channels where the source and destination share the same set of possible messages, so that the notion of “exact copy” is simple to define, our measures can be generalized to continuous-state channels as well as other possible definitions of copying. In Section IV, we show that our measure can be used to quantify the thermodynamic efficiency of physical copying processes, a central topic in the biological domain. In Section V, we demonstrate our measures on a real world dataset of amino acid substitution rates.

## II. COPY AND TRANSFORMATION INFORMATION

We first derive our proposed measure of copy information from four simple and intuitive axioms, and then construct the decomposition of mutual information into *copy* and *transformation*. Before proceeding, we briefly present some basic concepts from information theory that will be useful for our further developments.

### A. Preliminaries

We use the random variables *X* and *Y* to indicate the source and destination, respectively, of a communication channel (as defined in detail below). We assume that the source *X* and destination *Y* both take outcomes from the same countable set *𝒜*. We use Δ to indicate the set of all probability distributions whose support is equal to, or a subset of, *𝒜*. We use notation like *p*_*Y*_, *q*_*Y*_, …∈ Δ to indicate marginal distributions over *Y*, and *p*_*Y* |*x*_, *q*_*Y*| *x*_, …∈ Δ to indicate conditional distributions over *Y*, given the event *X* = *x*. Where clear from context, we will simply write *p*(*y*), *q*(*y*), …and *p*(*y* |*x*), *q*(*y* |*x*), …, and drop the subscripts.

For any two distributions *s* and *q* over the same set of outcomes, the *Kullback-Leibler* (KL) divergence is defined as

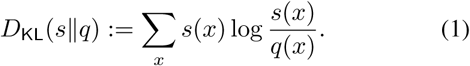

In this paper we will make use of the KL between Bernoulli distributions — that is, distributions over two states of the type (*a,* 1− *a*) — which is sometimes called “binary KL”. We will use the notation `d`(*a, b*) to indicate the binary KL,

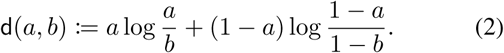

KL is always non-negative, and equal to 0 if and only if *s*(*x*) = *q*(*x*) for all *x*. Throughout this paper, we assume base 2 (meaning that information is measured in bits), unless otherwise noted.

In information theory, a *communication channel* specifies the conditional probability distribution of receiving different messages at a destination given messages transmitted by a source. Let *p*_*Y* |*X*_ (*y*|*x*) indicate such a conditional probability distribution. The amount of information transferred across a communication channel, given some probability distribution of source messages *s*_*X*_ (*x*), is quantified using the *mutual information* (MI) between the source and the destination [2],

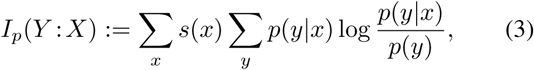

where *p*(*y*) is the marginal probability of receiving message *y* at the destination, defined as

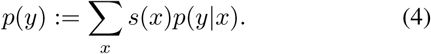

When writing *I*_*p*_(*Y* : *X*), we will omit the subscript *p* indicating the channel where it is clear from context. MI is a fundamental measure of information transmission, and can be operationalized in numerous ways [2]. It is non-negative, and large when (on average) the uncertainty about the message at the destination is low, given the source message.

Importantly, MI can also be written as a weighted sum of so-called *specific MI*[32] terms [33], one for each outcome of *X*,

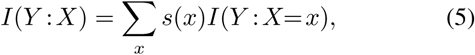

where the specific MI for outcome *x* is given by

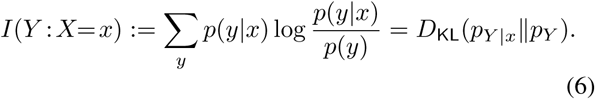

Each *I*(*Y* : *X* = *x*) indicates the contribution to MI arising from the particular source message *x*. We will sometimes use the term *total mutual information* (total MI) to refer to Eq. (3), so as to distinguish it from specific MI.

Specific MI also has an important Bayesian interpretation. Consider an agent who begins with a set of prior beliefs about *Y*, as specified by the prior distribution *p*_*Y*_ (*y*). Upon observing the event *X* = *x*, the agent updates their beliefs, resulting in the posterior distribution *p*_*Y* |*x*_(*y*). The KL divergence between the posterior and the prior, *D*_KL_(*p*_*Y* |*x*_ ‖*p*_*Y*_) (Eq. (6)), is called *Bayesian surprise* [34], and quantifies the amount of information acquired by the agent. It reaches its minimum value of zero, indicating that no information is acquired, if and only if the prior and posterior distributions match exactly. Bayesian surprise plays a fundamental role in Bayesian theory, including in the design of optimal experiments [35–38] and the selection of “non-informative priors” [39, 40]. Specific MI is a special case of Bayesian surprise, when the prior *p*_*Y*_ is the marginal distribution at the destination, as determined by a choice of source distribution *s*_*X*_ and channel *p*_*Y* |*X*_ according to Eq. (4). In general, however, Bayesian surprise may be defined for any desired prior *p*_*Y*_ and posterior distribution *p*_*Y | x*_, without necessarily making reference to a source distribution *s*_*X*_ and a communication channel *p*_*Y* |*X*_.

Because Bayesian surprise it is a general measure that includes specific MI as a special case, we will formulate our analysis of copy and transformation information in terms of *D*_KL_(*p*_*Y*| *x*_ *p*_*Y*_). Note that while the notation *p*_*Y* |*x*_ implies conditioning on the event *X* = *x*, formally *p*_*Y* |*x*_ can be any distribution whatsoever. Thus, we do not technically require that there exist some full joint or conditional probability distribution over *X* and *Y*. Throughout the paper we will refer to the distributions *p*_*Y* |*x*_ and *p*_*Y*_ as the “posterior” and “prior”.

Proofs and derivations are contained in the appendix.

### B. Axioms for copy information

To derive a measure of copy information, we propose four axioms for copy information. Our axioms are motivated by the following ideas: First, our decomposition should apply at the level of individual source message, i.e., we wish to be able to decompose each specific mutual information term (or more generally, Bayesian surprise) into a *(specific) copy information* term and a non-negative *(specific) transformation information* term. Second, we postulate that if there are two channels with the same marginal distribution at the destination, then the channel with the larger *p*(*x* |*x*) (probability of destination getting message *x* when the source transmits message should have larger copy information for source message *x* (this is, so to speak, our “central axiom”). Alternatively, under the Bayesian interpretation, this postulate can be interpreted as saying that if two Bayesian agents both start with the same prior and receive a piece of evidence *x*, then copy information should be greater for the agent whose posterior probability of *x* is larger.

Formally, we assume that each copy information term is a real-valued function of the posterior distribution, the prior distribution, and the source message *x*, written generically as *F* (*p*_*Y* |*x*_, *p*_*Y*_, *x*). Given any measure of copy information *F*, the transformation information associated with message *x* is then the remainder of *D*_KL_(*p*_*Y* |*x*_*‖p*_*Y*_) beyond *F*,

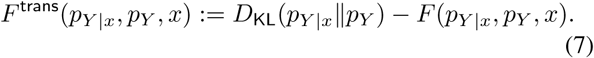

We now propose a set of axioms that any measure of copy information should satisfy.

First, we postulate that copy information should be bounded between 0 and the Bayesian surprise, *D*_KL_(*p*_*Y* |*x*_*Ip*_*Y*_). Given Eq. (7), this guarantees that both *F* and *F* ^trans^ are non-negative.

**Axiom 1.** *F* (*p*_*Y* |*x*_, *p*_*Y*_, *x*) *≥* 0.

**Axiom 2.** *F* (*p*_*Y* |*x*_, *p*_*Y*_, *x*) *≤ D*_KL_(*p*_*Y* |x_‖p_Y_).

Then, we postulate that copy information for source message *x* should increase monotonically as the posterior probability of *x* increases, assuming the prior distribution is held fixed. (This is the “central axiom” mentioned above.)

**Axiom 3.** *If p*_*Y* |*x*_(*x*) *≤ q*_*Y* |*x*_(*x*), *then F* (*p*_*Y* |*x*_, *p*_*Y*_, *x*) *≤F* (*p*_*Y* |*x*_, *p*_*Y*_, *x*)

In Appendix C, we show that any copy information that satisfies the above three axioms must obey *F* (*p*_*Y*| *x*_, *p*_*Y*_, *x*) = 0 whenever *p*_*Y* |*x*_(*x*) ≤ *p*_*Y*_ (*x*). What happens in the case *p*_*Y* |*x*_(*x*) *> p*_*Y*_ (*x*), however, is not totally determined. In fact, there are some trivial functions, such as *F* (*p*_*Y* |*x*_, *p*_*Y*_, *x*) = 0 for all *p*_*Y*| *x*_, *p*_*Y*_, and *x*, that satisfy the above axioms. Such trivial cases are excluded by our final axiom, which states that for all prior distribution and all posterior probabilities *p* (*x*) *> p*_*Y*_ (*x*), there are posterior distributions that contain *only* copy information.

**Axiom 4.** *For any p*_*Y*_ *and c* ∈ [*p*_*Y*_ (*x*), 1], *there exists a posterior distribution p*_*Y* |*x*_ *such that p*_*Y* |*x*_(*x*) = *c and F* (*p*_*Y* |*x*_, *p*_*Y*_, *x*) = *D*_KL_(*p*_*Y* |*x*_*Ip*_*Y*_).

### C. The measure 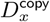

We now present the unique measure, which we call 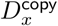, that satisfies the copy information axioms proposed in the last section. Given a prior distribution *p*_*Y*_, posterior distribution *p*_*Y* |*x*_, and source message *x*, 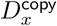 is defined as

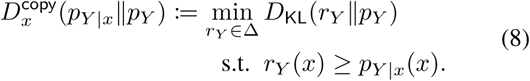

The minimization problem of Eq. (8) turns out to have a simple closed-form solution, as shown in Appendix A:

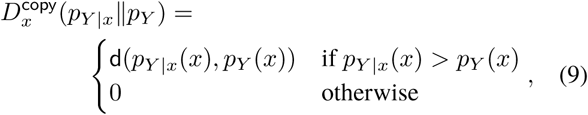

where we have used the notation of Eq. (2). It turns out that 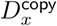 is the *unique* measure that satisfies the four proposed axioms. We can now state the main result of our paper:

#### Theorem 1.

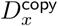 *is the unique measure which satisfies Axioms 1 to 4.*

In the Appendix B we demonstrate that 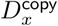 satisfies all the axioms, and in the Appendix C we prove that it is actually the only measure that satisfies them. We further show that if we drop Axiom 4, then 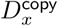 is the largest possible measure that can satisfy the remaining axioms. Finally, note that given the definition of *F* ^trans^ in Eq. (7), 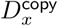 also defines a particular, non-negative measure of transformation information, which we call 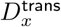,

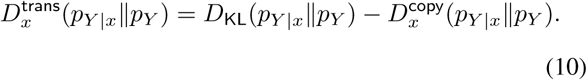

### D. Decomposing mutual information

We now show that 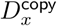 and 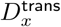 provide the ingredients for a decomposition of mutual information (MI) into *MI due to copying* and *MI due to transformation*. Recall that MI can be written as an expectation over specific MI terms, as shown in Eq. (6). Each specific MI term can be seen as a Bayesian surprise, where the prior distribution is the marginal distribution at the destination (see Eq. (4)), and the posterior distribution is the conditional distribution of destination given a particular source message under the channel. Thus, our definitions of 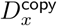 and 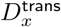 provide a non-negative decomposition of each specific MI term,

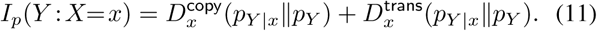

In consequence, they also provide a non-negative decomposition of the total MI into two non-negative terms: the *total copy information* and the *total transformation information*,

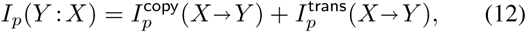

where 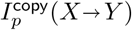 and 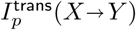 are given by

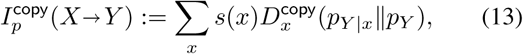

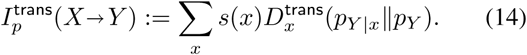

(When writing *I*^copy^ and *I*^trans^, we will often omit the subscript *p* where the channel is clear from context.)

In the definitions of 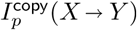 and 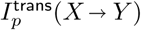, the notation *X* → *Y* indicates that *I*^copy^ and *I* are defined as the copy and transformation information when *X* is the source and *Y* is the destination. This is necessary because, unlike MI, *I*^copy^ and *I*^trans^ are in general non-symmetric, so it is possible that *I*^copy^(*X* →*Y*) */*= *II*^copy^ (*Y* → *X*), and similarly for *I*^trans^. We also note that the above form of *I*^copy^ and *I*^trans^, where they are written as sums over individual source message, is sometimes referred to as *trace-like* form in the literature, and is a common desired characteristic of information-theoretic functionals [41, 42].

For illustration purposes, in Fig. 2 we plot the behavior of *I*^copy^ and *I*^trans^ in the classical binary symmetric channel (BSC) (see caption for details). More detailed analysis of copy and transformation informatoin in the BSC is discussed in Appendix F.

**Figure 2.**
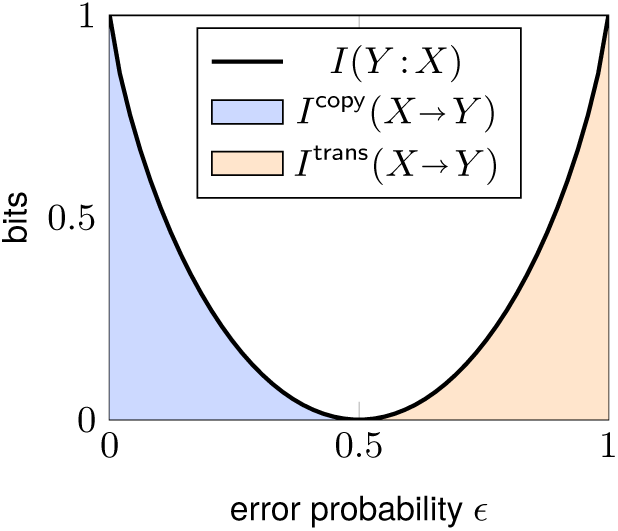
The Binary Symmetric Channel (BSC) with a uniform source distribution. Values of mutual information, *I*^copy^(*X* →*Y*) (Eq. (13)) and *I*^trans^(*X* →*Y*) (Eq. (14)) for the BSC along the whole range of error probabilities *ϵ* ∈ [0, 1]. When *ϵ ≤* 1*/*2, all mutual information is *I*^copy^ (blue shading), while when *ϵ ≥* 1*/*2, all mutual information is *I*^trans^ (red shading).

Finally, we note that by a simple manipulation, we can decompose the marginal the entropy of the destination *H*(*Y*) into three non-negative components:

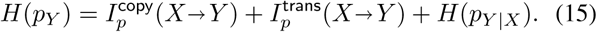

Thus, given a channel from *X* to *Y*, the uncertainty in *Y* can be written as the sum of the copy information from *X*, the transformed information from *X*, and the intrinsic noise in that channel from *X* to *Y*.

### E. Copying efficiency

As mentioned in the Introduction, our approach provides a way to quantify which portion of the information transmitted across a channel is due to copying rather than transformation, which we refer to as “copying efficiency”. Copying efficiency is defined at the level of individual source messages as

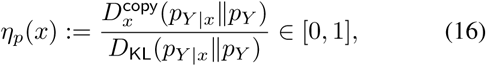

where the bounds come directly from Axioms 1 and 2. It can also be defined at the level of a channel as whole as

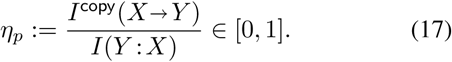

The bounds follow simply given the above results.

For Eq. (16) and Eq. (17) to be useful efficiency measures, there should exist channels which are either “completely inefficient” (have efficiency 0) or “maximally efficient” (achieve efficiency 1). For the case of Eq. (16), the bounds can be saturated because of Axiom 4, which guarantees that for any source message *x*, prior *p*_*Y*_, and desired posterior probability *p*_*Y* |*x*_(*x*) *> p*_*Y*_ (*x*), there exists a posterior *p*_*Y*| *x*_ such that the Bayesian surprise *D*_KL_(*p*_*Y*_ |_*x*_ ‖ *p*_*Y*_) is composed entirely of copy information (for example, see Eq. (B1)).

One can show that the bounds in Eq. (17) can also be saturated. First, it can be verified that completely inefficient channels exists, since any channel which has *p*_*Y* |*x*_(*x*) *≤ p_Y_* (*x*) for all *x* ∈*𝒜* will have *I*^copy^(*X*→ *Y*) = 0 (note that such channels exists at all levels of mutual information). We also prove that maximally efficient channels exist, using the following result which is proved in Appendix E.

#### Proposition 1.

*For any source distribution s_X_ with full support and H*(*s*_*X*_) *< ∞, there exist channels p for all levels of mutual information I*_*p*_(*Y* : *X*) ∈ [0, *H*(*s*_*X*_)] *such that* 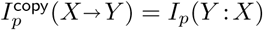.

## III. GENERALIZATION AND RELATION TO RATE DISTORTION

In this section, we show that 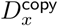 can be written as a particular element among a broad family of copy information measures, which generalize the formal definition of what is meant by “copying”. All elements of this family of copying information measures have the following form:

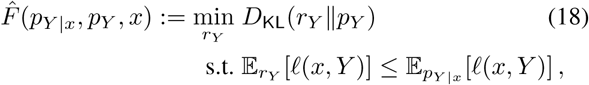

where *ℓ* (*x, y*) is some function that specifies the “loss” the occurs when source message *x* is received by the destination as message *y*. 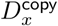 has the form of Eq. (18) if one takes *ℓ* (*x, y*) := 1 − *δ*(*x, y*), which is called “0-1 loss” in statistics [43] and “Hamming distortion” in information theory [2], and which formalizes copy information in terms of exact identity between source and destination message.

Each particular loss function induces its own measure of copy information, allowing us to define copy information in a more general manner. While 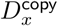, as defined via the 0-1 loss function, is a simple and reasonable choice in a variety of applications, in some cases copy information could also be defined in a more general way, in terms of another loss function. One important application of this generalization is the situation where copying is not defined in terms of simple identity between source and destination messages, but rather in terms of some more continuous measure of similarity. This is especially useful for continuous-valued random variables, because in this case the constraint in Eq. (8) is no longer meaningful (it only constraints a measure-0 property of *r*_*Y*_). Here a natural measure of copy information is given by the squared error loss function, *ℓ* (*x, y*) := (*x* − *y*)^2^, giving

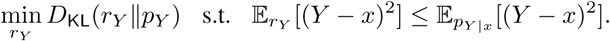

Note that this optimization problem has been investigated in the maximum entropy literature, and has been shown to be particularly tractable when *p*_*Y*_ belongs to an exponential family [44–46].

Recall that we have so far assumed that the source and destination share the same set of possible outcomes, *𝒜*. Importantly, with a generalized copy information the case where the source and destination do not share the same set of outcomes can be studied. In that case, some measure of similarity, must be defined between source and destination messages.

The optimization problem of Eq. (18) has some interesting properties. First, it is an example of a “minimum cross-entropy” distribution, which itself is a generalization of the “maximum entropy” principle [47, 48]. Using this fact, in Appendix D we show that the optimal solution has the following simple form

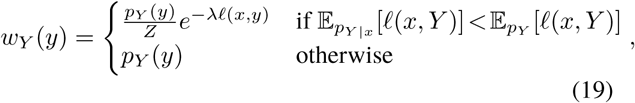

where *Z* is an appropriate normalization constant and *λ >* 0 is a Lagrange multiplier chosen so that the constraint Eq. (18) is satisfied. Thus, Eq. (18) can be solved efficiently by searching across the 1-dimensional space of possible *λ* values.

In Appendix D, we also show that Eq. (18) emerges as the unique measure that satisfies our aforementioned s et of four axioms, as long as Axiom 3 and Axiom 4 are replaced by their “generalized” versions. The generalized version of Axiom 3 states that one posterior distribution should have higher copy information than another whenever its expected loss is lower:

**Axiom 3***\*. If* 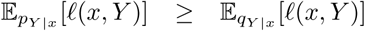, *then F* (*p*_*Y* |*x*_, *p*_*Y*_, *x*) *≤ F* (*q*_*Y* |*x*_, *p*_*Y*_, *x*).

The generalized version of Axiom 4 states that at all expected loss values lower than that achieved by *p*_Y_, there are channels which transmit information only by occupying.

**Axiom 4***\*. For any p*_*Y*_ *and* 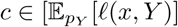, min_*y*_ ℓ (*x, y*)], there exists a posterior distribution p_*Y* |*x*_ *such that* 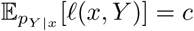 *and F* (*p*_*Y* |*x*_, *p*_*Y*_, *x*) = *D*_KL_(*p*_*Y* |x_‖p_Y_).

Note that we used that min_*y*_ *ℓ* (*x, y*)] is the lowest possible expected loss that can be achieved by any distribution.

We finish by noting that Eq. (18) shows a similarity between our approach and so called *rate-distortion theory* [2]. In rate distortion theory, one is given a distribution over source messages *s*_*X*_ and a “distortion function” *ℓ* (*x, y*) which specifies the loss incurred when source message *x* is encoded with destination message *y*. The problem is to find the channel *p*_*Y* |*X*_ which minimizes mutual information without exceeding some constraint on the expected distortion,

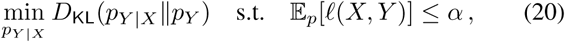

where *α* is an externally-determined parameter. The prototypical application of rate-distortion theory is compression, i.e., finding representations of the source that are both compressed (have low mutual information) and have low distortion. As can be seen by comparing Eq. (18) and Eq. (20), the optimization problem considered in our definition of copy information and the optimization found in rate distortion are quite similar: they both involve minimizing a KL divergence subject to an expected loss constraint. Nonetheless, there are some important differences. First and foremost, the goals of the two approaches are different: we aim to decompose the information transmitted by a fixed externally-specified channel into copy and transformation (it is this given channel that determines the constraint *α* in Eq. (18)), while in rate-distortion there is no externally-specified channel and the aim is instead to find an optimal channel *de novo*. In this respect, our approach also has some similarities to the so-called *minimum information principle*, which aims to quantify how much information about sent messages is carried by different properties of received messages [49]. Second, our approach is motivated by a set of axioms which postulate how a measure of copy information should behave, rather than from channel-coding considerations as is used to derive the optimization problem in rate-distortion [2]. Lastly, copy information is defined in a point-wise manner for each source message *x*, rather than for an entire set of source messages at once as is rate-distortion. However, we do note that in principle it should be possible to define Eq. (18) in a way that include copy information measures defined in a channel-wise manner (like Eq. (20)) rather than a pointwise manner (like Eq. (8)).

## IV. THERMODYNAMIC COSTS OF COPYING

Given the close connection between information theory and statistical physics, many information-theoretic quantities can be interpreted in thermodynamic terms [8]. As we show here, this includes our proposed measure of copy information,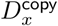. Specifically, we will show that 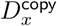 reflects the minimal amount of thermodynamic work necessary to copy a physical entity such as a polymer molecule. This latter example emphasizes the important difference between information transfer by copying versus by transformation in a fundamental, biologically-inspired physical setup.

Consider a physical system coupled to a heat bath at temperature *T*, and which is initially in equilibrium distribution 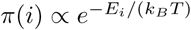 with respect to some Hamiltonian *E* (*k*_*B*_ is Boltzmann’s constant). Now imagine that the system is driven to some non-equilibrium distribution *p* by a physical process, and that by the end of the process the Hamiltonian is again equal to *E*. The minimal amount of work required by any such process is related to the KL divergence between *p* and *π* [50],

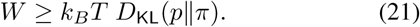

The limit is achieved by thermodynamically-reversible processes. (In this subsection, in accordance with the convention in physics, we assume that all logarithms are in base *e*, so information is measured in nats.)

Recent work has analyzed the fundamental thermodynamic constraints on copying in a physical system, for example for an information-carrying polymer like DNA [51, 52]. Here we will generally follow the model described in [51], while using our notation and omitting some details that are irrelevant for our purposes (such as the microstate/macrostate distinction). In this model, the source *X* represents the state of the original system (e.g., the sequence of the polymer to be copied), and the destination *Y* represents the state of the product (e.g., the sequence that is produced by the copying mechanism). We imagine that the original system is in state *X* = *x*, and that replicates are produced according to the conditional distribution *p*_*Y*|*x*_(*x*). Finally, assume that *X* and *Y* are statistically independent under the equilibrium distribution (in other words, that there the energetic coupling between *X* and *Y* is negligible compared to their internal energies), and let *π*_*Y*_ indicate the equilibrium distribution of the replicate *Y*. Following Eq. (21), the minimal work required to bring system *Y* out of equilibrium and produce replicates of *x* according to *p*_*Y*|*x*_(*x*) is given by

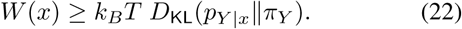

Note that Eq. (22) specifies the minimal work required to create the overall distribution *p*_*Y* |*x*_. However, in many real-world scenarios, likely including DNA copying, the primary goal is to create exact copies of the original state, not transformed versions it (such as nonrandom mutations). That means that for a given source state *x*, the quality of the replication process can be quantified by the probability of making an exact copy, *p*_*Y* |*x*_(*x*). We can now ask: what is the minimal work required by any physical replication process whose probability of making exact copies is at least as large as *p*_*Y* |*x*_(*x*)? To make the comparison fair, we require that the process operate under the same equilibrium distribution, *π*_*Y*_. The answer is given by the minimum of the RHS of Eq. (22) under a constraint on the exact-copy yield, which is exactly proportional to 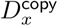:

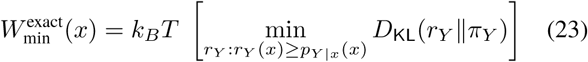

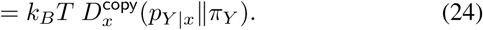

The additional work that is expended by the replication process is then lower bounded by a quantity proportional 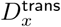,

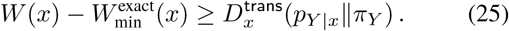

This shows formally the intuitive idea that transformation information contributes to thermodynamic costs but not to the accuracy of correct copying.

In most cases, a replication system is designed for copying not just one source state *x*, but an entire ensemble of source states, which can be assumed to be distributed according to some *s*_*X*_ (*x*) (for example, the DNA replication system can copy a huge of ensemble of source DNA sequences, not just one). Across the entire ensemble of source states, the minimal amount of *expected* thermodynamic work required to produce replicates according to conditional distribution *p*_*Y* |*X*_ is given by

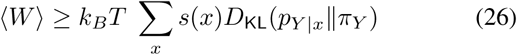

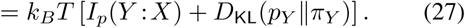

Since KL is non-negative, the minimum expected work is lowest when the equilibrium distribution *π*_*Y*_ matches the marginal distribution of replicates, *p*_*Y*_ (*y*) =∑_x_ *s*(*x*)*p*(*y*|*x*). Using similar arguments as above, we can ask about the minimum expected work required to produce replicates, assuming each source state *x* achieves an exact-copy yield of at least *p*_*Y* |*x*_(*x*). This turns out to be the expectation of Eq. (24),

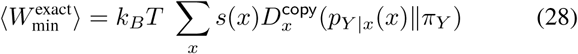

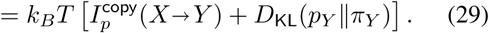

The additional expected work that is needed by the replication process, above and beyond an optimal process that achieves the same exact-copy yield, is lower bounded by the transformation information,

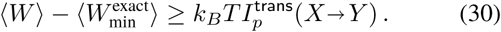

When the equilibrium distribution *π*_*Y*_ matches the marginal distribution *p*_*Y*_, 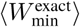 is exactly equal *k*_*B*_*TI*^copy^. Further-more, in this special case the thermodynamic efficiency of exact copying, defined as the ratio of minimal work to actual work, becomes equal to the information-theoretic copying efficiency of *p*, as defined in Eq. (17):

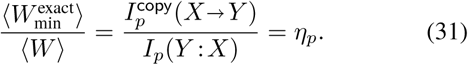

As can be seen, standard information-theoretic measures, such as Eq. (22), bound the minimal thermodynamic costs of transferring information from one physical system to another, whether that transfer happens by copying or by transformation. However, as we have argued above, the difference between copying and transformation is essential in many biological scenarios, as well as other domains. In such cases, 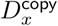 arises naturally as the minimal thermodynamic work required to replicate information by copying. Concerning the example of the DNA copying that we followed throughout this section, we note that these results should be interpreted with some care. We have generally imagined that the source system represents the state of an entire polymer, e.g., the state of an entire DNA molecule, and that the probability of exact copying refers to the probability that the entire sequence is reproduced without any errors. Alternatively, one can use the same framework to consider probability of copying a single monomer in a long polymer (assuming that the thermodynamics of polymerization can be disregarded), as might be represented for instance by a single-nucleotide DNA substitution matrix [53], as analyzed in the last section. Generally speaking, 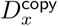 computed at the level of single monomers will be different from 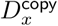 computed at the level of entire polymers, since the probability of exact copying means different things in these two formulations.

## V. COPY AND TRANSFORMATION IN AMINO ACID SUBSTITUTION MATRICES

In the previous section, we saw how 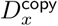 and *I*^copy^ arises naturally when studying the fundamental limits on the thermodynamics of copying, which includes the special case of replicating information-bearing polymers. Here we demonstrate how these measures can be used to characterize the information-transmission properties of a real-world biological replication system, as formalized by a communication channel *p*_*Y*|*X*_ from parent to offspring [53, 54]. In this context, we show how *I*^copy^ can be used to quantify precisely how much information is transmitted by copying, without mutations. At the same time, we will use *I*^trans^ to quantify how much information is transmitted by transformation, that is by systematic *nonrandom* mutations that carry information but do not preserve the identity of the original message [16, 17]. We also quantify the effect of purely-random mutations, which correspond to the conditional entropy of the channel, *H*(*p*_*Y*|*X*_).

We demonstrate these measures on empirical data of *point accepted mutations* (PAM) of amino acids. PAM data represents the rates of substitutions between different pairs of amino acids during the course of biological evolution, and has various applications, including evolutionary modeling, phylogenetic reconstructions, and protein alignment [54]. We emphasize that PAM matrices between amino acids do not reflect the direct physical transfer of information from protein to protein, but rather the result effects of underlying processes of DNA-based replication and selection, followed by translation.

Formally, a PAM matrix *Q* for amino acids is a continuous-time rate matrix, where *Q*_*yx*_ represents the instantaneous rates of substitutions from amino acid *x* to amino acid *y*, where both *x* and *y* belong to 𝒜 = {1, *…,* 20}, representing the 20 standard amino acids. We performed our analysis on a fixed PAM matrix *Q*, which was published by Le and Gascuel [54] (this matrix was provided by the `pyvolve` Python package [55]). We calculated a discrete-time conditional probability distribution *p*_*Y*|*X*_ from this matrix by taking the matrix exponential, *p*_*Y*|*X*_ = exp(*τQ*). Thus, *p*(*y* |*x*) represents the probability that amino acid *x* is replaced by amino acid *y* over time scale *τ*. For simplicity, we used timescale *τ* = 1 and assumed that the source distribution over amino acids is uniform, *s*(*x*) = 1*/*20 for all *x*. Using the decomposition presented in Eq. (14), we get the following values for the communication channel described by the conditional probabilities *p*_*Y*|*X*_ :

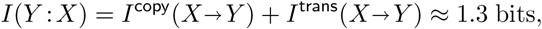

Where

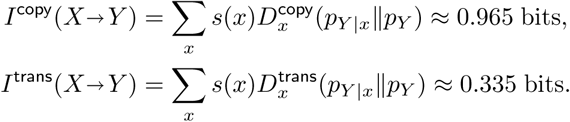

We also computed the intrinsic noise for this channel (see Eq. (15)),

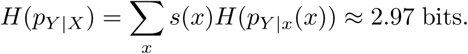

Finally, we computed the specific copy and transformation information, 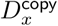 and 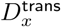, for different amino acids. The results for different amino acids are shown in Fig. 3. We remind the reader that the sum of 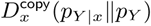 and 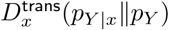 for each amino acid *x* — that is, the total height of the stacked bar plots in the figure — is equal to the specific MI *I*(*Y* : *X* = *x*) for that *x*, as explained in the decomposition shown in Eq. (11).

**Figure 3.**
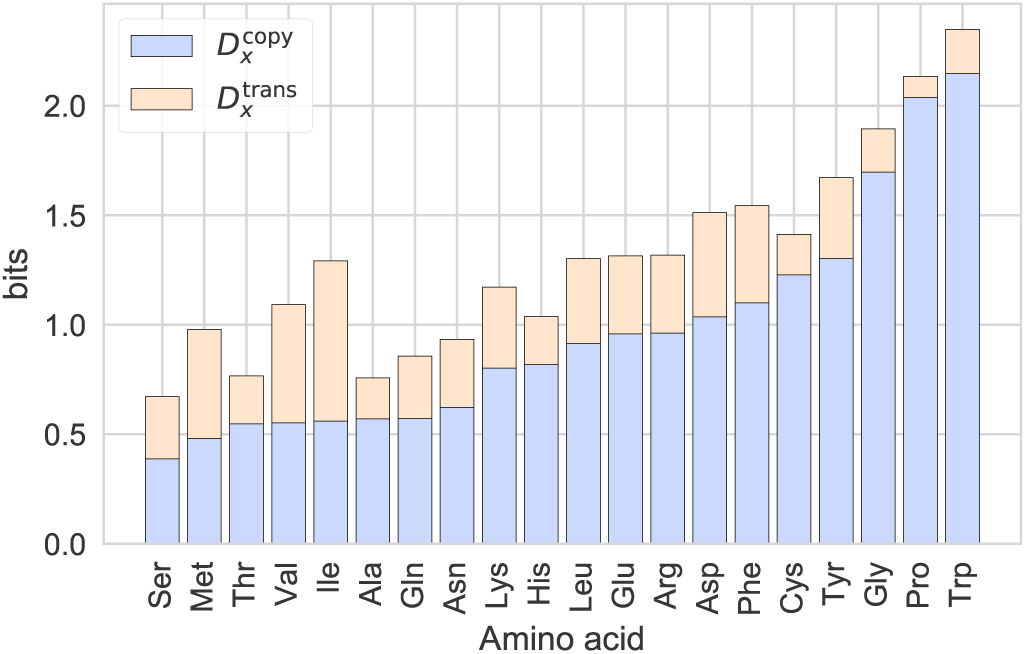
Copy and transformation information for different amino acids, based on an empirical PAM matrix [54]. In blue, we show magnitude of 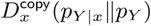, on top, the amount of transformed information, in red. The sum of both is the specific MI for each amino acid, according to the decomposition given in equation (11)

While we do not dive deeply in the biological significant of these results, we highlight several interesting findings. First, for this PAM matrix and timescale (*τ* = 1), a considerable fraction of the information (≈ 1*/*3) is transmitted not by copying but by non-random mutations. Generally, such non-random mutations represent underlying physical, genetic, and biological constraints that allow some pairs of amino acids to substitute each other more readily than other pairs.

Second, we observe considerable variation in the amount of specific MI, copy information, and transformation between different amino acids, as well as different the ratio of copy information to transformation information. In general, amino acids with more copy information are conserved unchanged over evolutionary timescales. At the same time, it is known that conserved amino acids tend to be “outliers” in terms of their physiochemical properties (such as hydrophobicity, volume, polarity, etc.), since mutations to such outliers are likely to alter protein function in deleterious ways [56, 57]. To analyze this quantitatively, we used Miyata’s measure of distance between amino acids, which is based on differences in volume and polarity [58]. For each amino acid, we quantified its degree of “outlierness” in terms of its mean Miyata distance to all 19 other amino acids. The Spearman rank correlation between this outlierness measure and copy information (as shown in Fig. 3) was 0.66 (*p* = 0.0035). On the other hand, the rank correlation between outlierness and transformation information was 0.02 (*p* = 0.92). Similar results were observed for other chemically-motivated measures of amino acid distance, such as Grantham’s distance [59] and Sneath’s index [60]. This demonstrates that amino acids with unique chemical characteristics tend to have more copy information, but not more transformation information.

## VI. DISCUSSION

Although mutual information is a very common and successful measure of transmitted information, it is insensitive to the distinction between information that is transmitted by copying versus information that is transmitted by transformation. Nonetheless, as we have argued, this distinction is of fundamental importance in many real-world systems.

In this paper we propose a rigorous and practical way to decompose specific mutual information, and more generally Bayesian surprise, into two non-negative terms corresponding to copy and transformation, *I* = *I*^copy^ + *I*^trans^. We derive our decomposition using an axiomatic framework: we propose a set of four axioms any measure of copy information should obey, and then identify the unique measure that satisfies those axioms. At the same time, we show that our measure of copy information is one of a family of functionals, each of which corresponds to a different way of quantifying error in transmission. We also demonstrate that our measures have a natural interpretation in thermodynamic terms, which suggests novel approaches for understanding the thermodynamic efficiency of biological replication processes, in particular DNA and RNA duplication. Finally, we demonstrate our results on real-world biological data, exploring copy and transformation information of amino acid substitution rates. We find significant variation among the amount of information transmitted by copying vs. transformation among different amino acids.

Several directions for future work present themselves. First, there is a large range of practical and theoretical application of our measures, ranging form analysis of biological and neural information transmission to the study of the thermodynamics of self-replication, a fundamental and challenging problem in biophysics [61]. Second, we suspect our measures of copy and transformation information have further connections to existing formal treatments in information theory, in particular rate-distortion theory [2], whose connections we started to explore here. We also believe that our decomposition may be generalizable beyond Bayesian surprise and mutual information to include other information-theoretic measures, including conditional mutual information and multi-information. Decomposing conditional mutual information is of particular interest, since it will permit a decomposition of the commonly-used *transfer entropy* [62] measure into copy and transformation components, thus separating two different modes of information flow between systems. Finally, our decomposition of mutual information into copy and transformation components has some high-level similarities to other recent proposals for non-negative decompositions of information measures, for example non-negative decompositions of multivariate information into redundant and synergistic components [63], integrated information decompositions [64, 65], and decompositions of mutual information into “semantic” (valuable) and “non-semantic” (non-valuable) information [66]. Further research should explore if and how our decomposition into copy and transformation information relates to these existing approaches.

## ACKNOWLEDGMENTS

AK was supported by Grant No. FQXi-RFP-1622 from the FQXi foundation, and Grant No. CHE-1648973 from the U.S. National Science Foundation. AK would like to thank the Santa Fe Institute for supporting this research. The authors thank Jordi Fortuny, Rudolf Hanel, Joshua Garland, and Blai Vidiella for helpful discussions on the manuscript.

## Appendix A: Derivation of Eq. 9

Consider the optimization problem:

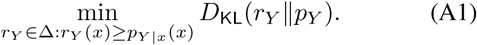

It is clear that if *p*_*Y*|*x*_(*x*) ≤ *p*_*Y*_(*x*), then the solution *r*_*Y*_(*y*) = *p*_*Y*_ (*y*) for all *y* satisfies the constraint, and achieves *D*_KL_(*p*_*Y*_‖*p*_*Y*_) = 0, the minimum possible. We now consider the case when *p*_*Y*|*x*_(*x*) > *p*_*Y*_(*x*). In this case, we use the chain rule for KL divergence [2] to write

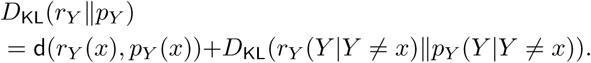

The second term is minimized by setting *r*_*Y*_(*y*) ∝ *p*_*Y*_(*y*) for *y* ≠ *x*, so that *r*_*Y*_ (*y*|*Y* ≠ *x*) = *p*_*Y*_ (*y*|*Y* ≠ *x*) and *D*_KL_(*r*_*Y*_ (*Y*|*Y* ≠ *x*)‖ *p*_*Y*_ (*Y*|*Y* ≠ *x*)) = 0. Thus, in the case that *p*_*Y*|*x*_(*x*) > *p*_*Y*_ (*x*), we have reduced the optimization problem of Eq. (A1) to the equivalent problem

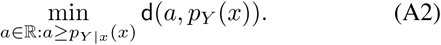

Note that the derivative `d`(*a, b*) with respect to *a* is 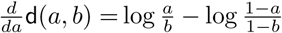, which is strictly positive when *a* > *b*. Given the assumption that *p*_*Y*|*x*_(*x*) > *p*_*Y*_(*x*), it must be that Eq. (A2) is minimized by setting *a* = *p*_*Y*|*x*_(*x*), the minimum possible. Thus, `d`(*p*_*Y*|*x*_(*x*), *p*_*Y*_(*x*)) is the solution to Eq. (A1) when *p*_*Y*|*x*_(*x*) > *p*_*Y*_ (*x*).

## Appendix B: 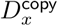 satisfies the four axioms

It satisfies Axiom 1 by non-negativity of KL. It satisfies Axiom 2 when *p*_*Y*|*x*_(*x*) > *p*_*Y*_(*x*) because `d`(*p*_*Y*|*x*_(*x*), *p*_*Y*_ (*x*)) ≤ *D*_KL_(*p*_*Y*|*x*_‖*p*_*Y*_) by the data processing inequality for KL divergence [67, Lemma 3.11]; otherwise, when *p*_*Y*|*x*_ ≤ *p*_*Y*_ (*x*), 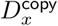 vanishes and thus satisfies Axiom 2 trivially. It satisfies Axiom 3 because if *p*_*Y*_ (*x*) ≤ *q*_*Y*_ (*x*), then the minimization problem of Eq. (8) for *q*_*Y*_ is more constrained than for *p*_*Y*_, so the minimum of the former must be larger than the minimum of the latter. Finally, to show that 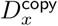 satisfies Axiom 4, for any prior distribution *pY* consider the following posterior distribution 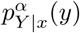:

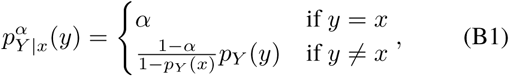

where *α* is a parameter that can vary from 0 to 1. It is easy to verify that for all *α* ∈ [*p*_*Y*_ (*x*), 1],

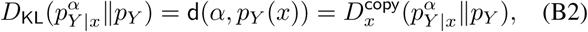

and that 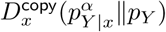 ranges in a continuous manner from 0 (for *α* = *p*_*Y*_(*x*)) to − log *p*_*Y*_(*x*) (for *α* = 1).

## Appendix C: Proof of Theorem 1

Before proceeding, we first prove two useful lemmas.

### Lemma C.1.

*Given Axiom 3, F* (*p*_*Y*|*x*_, *p*_*Y*_, *x*) = *F* (*q*_*Y*|*x*_, *p*_*Y*_, *x*) *if p*_*Y*|*x*_(*x*) = *q*_*Y*|*x*_(*x*).

### Proof.

Follows from applying Axiom 3 twice.

### Lemma C.2.

*Given Axioms 1 to 3, if p*_*Y*|*x*_(*x*) ≤ *p*_*Y*_ (*x*), *then F* (*p*_*Y*|*x*_, *p*_*Y*_,*x*) = 0.

### Proof.

If *p*_*Y*|*x*_(*x*) ≤ *p*_*Y*_ (*x*), then *F* (*p*_*Y*|*x*_, *p*_*Y*_, *x*) ≤ *F* (*p*_*Y*_, *p*_*Y*_, *x*) by Axiom 3. By Axiom 2, *F* (*p*_*Y*_, *p*_*Y*_, *x*) ≤ *D*_KL_(*p*_*Y*_‖*p*_*Y*_) = 0. Combining gives *F* (*p*_*Y*|*x*_, *p*_*Y*_,*x*) ≤ 0, while *F* (*p*_*Y*|*x*_, *p*_*Y*_, *x*) ≥ 0 by Axiom 1.

We then show that 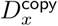 is the largest possible measure that satisfies Axioms 1 to 3.

### Proposition C.1.

*Any F which satisfies Axioms 1 to 3 must obey* 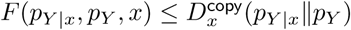.

### Proof.

Given Lemma C.2, without loss of generality we restrict our attention to the case where *p*_*Y*|*x*_(*x*) > *p*_*Y*_(*x*). Define the posterior 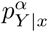 as in Eq. (B1), while taking *α* = *p*_*Y*|*x*_(*x*). Then, by Lemma C.1,

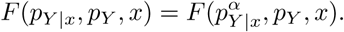

At the same time,

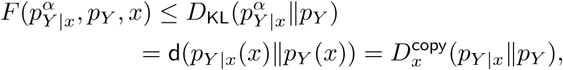

where the first inequality follows from Axiom 2, and the second equality from Eq. (B2).

We are now ready to prove the main result from Section II B.

### Proof of Theorem 1.

Consider some *p*_*Y*|*x*_, *p*_*Y*_, *x*, and assume *p*_*Y*|*x*_(*x*) > *p*_*Y*_ (*x*) (without loss of generality by Lemma C.2). By Axiom 4, there must exist a posterior *q*_*Y*|*x*_ such that *q*_*Y*|*x*_(*x*) = *p*_*Y*|*x*_(*x*) and

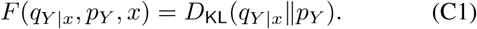

Note that by definition of 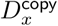 (and in consistency with Axiom 2), 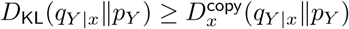.

Then, by Lemma C.1, *F* (*p*_*Y*|*x*_, *p*_*Y*_, *x*) = *F* (*q*_*Y*|*x*_, *p*_*Y*_, *x*) since *p*_*Y*|*x*_(*x*) = *q*_*Y*|*x*_(*x*). Similarly, it can be verified that 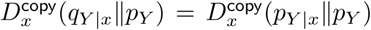. Combining the above results shows that 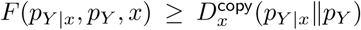. the theorem follows by combining with Proposition C.1.

## Appendix D: Solution and axiomatic derivation of Eq. 18

In this appendix, we first show that Eq. (19) is the solution to Eq. (18), the generalized copy information measure considered in Section III.

First, note that if 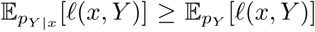, then the solution *r*_*Y*_ = *p*_*Y*_ satisfies the constraint and achieves the smallest possible KL value of 0. Therefore, without loss of generality, we will consider the case when 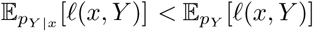. From the MaxEnt literature [48], it is known that the solution to the following equality-constrained optimization problem,

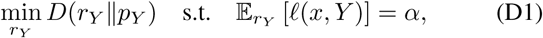

is given by

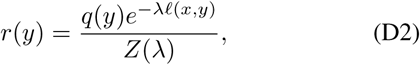

where *Z*(*λ*) is the normalization constant and *λ* is a Lagrange multiplier that enforces the constraint. This can be seen as a 1-dimensional manifold of distributions parameterized by *λ*. We show that as *λ* increases, 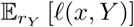 monotonically decreases while *D*(*r*_*Y*_‖*p*_*Y*_) monotonically increases. That means that solutions to the inequality-constrained optimization problem, Eq. (18), will make the constraint 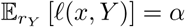 tight, and thus have the form of Eq. (D2). This recovers Eq. (19).

To show that 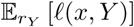 monotonically decreases while *D*(*r*_*Y*_‖*p*_*Y*_) monotonically increases with *λ*, we first write the derivatives of *Z*(*λ*) and *r*(*y*) with respect to *λ*,

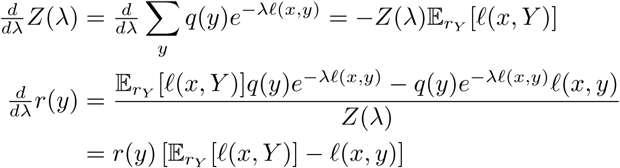

Clearly, the expectation of *f*(*x, y*) decreases with *λ*,

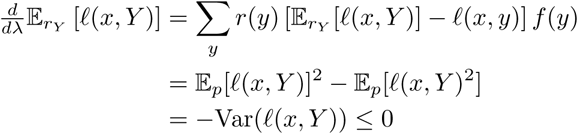

At this point, note that *r*_*Y*_ = *p*_*Y*_ when *λ* = 0. Since we are considering the case where 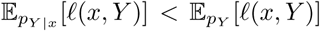, it must be that *λ* > 0. We can then show that the KL term increases with *λ*,

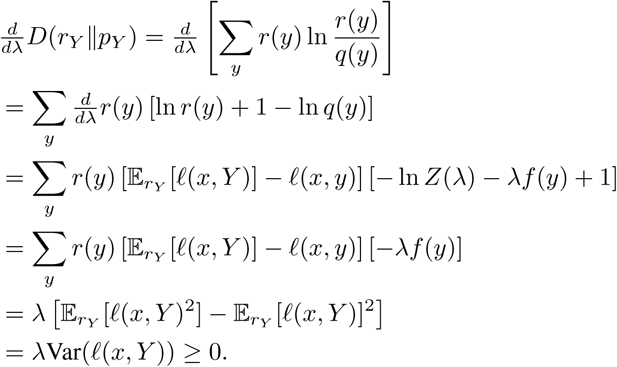

We now demonstrate that the generalized copy information defined in Eq. (18), 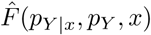, is the unique measure that satisfies Axioms 1 and 2 and our modified Axioms 3* and 4*. Our derivation has the same structure as the one in Appendix C, and we proceed more quickly.

First, by applying applying Axiom 3* twice, it is clear that

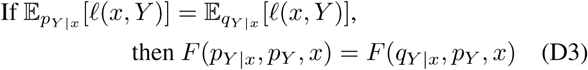

In addition, by Axioms 1 and 2, 0 ≤ *F* (*p*_*Y*_, *p*_*Y*_, *x*) ≤*D*_KL_(*p*_*Y*_‖*p*_*Y*_) = 0, so it must be that *F* (*p*_*Y*_, *p*_*Y*_, *x*) = 0. By Axiom 3*, that means that *F* (*p*_*Y*|*x*_, *p*_*Y*_, *x*) = 0 for any *p*_*Y*|*x*_ with 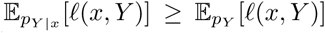 (i.e., whose loss is bigger than that of the prior distribution). In the rest of this appendix, we will assume without loss of generality that 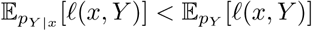.

We now show that 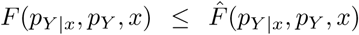. Given a choice of *p*_*Y*| *x*_, *p*_*Y*_, and *x*, let *w*_*Y*_ be the solution to Eq. (18), so

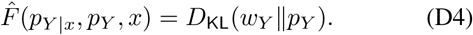

Given the derivations at the beginning of this appendix, we know that 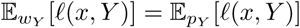, which implies

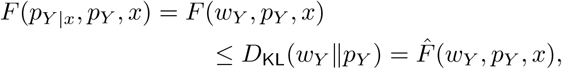

where the first e quality uses Eq. (D3), the inequality uses Axiom 2, and the last equality uses Eq. (D4).

Finally, we show that 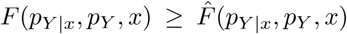. By Axiom 4*, there must exist a posterior *q*_*Y*|*x*_ such that 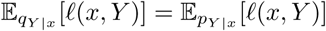 and

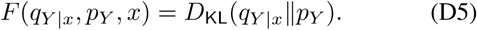

Note that by definition of 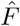 (and in consistency with Axiom 2), 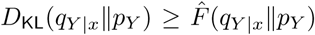. Then, by Eq. (D3), 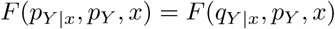. Similarly, it can be verified that 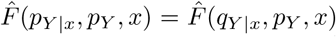. Combining the above results shows that 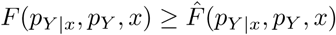.

Thus, 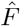 is the only function that satisfies Axioms 1 and 2 and our modified Axioms 3* and 4*

## Appendix E: Proof of Prop. 1

Before proving Proposition 1, we prove several intermediate results.

We start with a lemma that derives some useful properties of the roots of the quadratic polynomial *ax*^2^ − (*a* + *s*)*x* + *sc*.

### Lemma E.1.

*Consider the function*

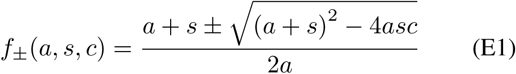

*where a* ∈ ℝ, *s* ∈ (0, 1], *c* ∈ [0, 1]. *Then,*

1. *If a* < 0, *f*_+_(*a, s, c*) ≤ 0 *and the inequality is strict when c* > 0. *Otherwise, when a ≥* 0, *f*_+_(*a, s, c*) ≥ 1, *and the inequality is strict when a* = 0.
2. 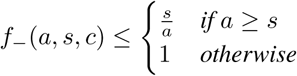.
3. 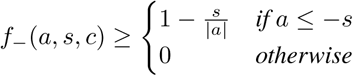.
4. lim_*a*→0_ *f*_−_(*a, s, c*) = *c.*
5. 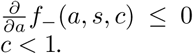, *and the inequality is strict when c* < 1.

### Proof. Proof of 1.

First consider *a* < 0, in which case *f*_+_(*a, s, c*) ≤ *f*_−_(*a, s, c*). Vieta’s formula states that

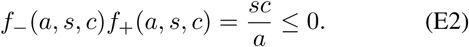

This implies that *f*_+_(*a, s, c*) must be less than 0. The inequality is strict when *c* > 0.

In the case *a* ≥ 0, we lower bound the determinant,

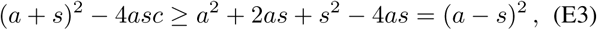

This implies

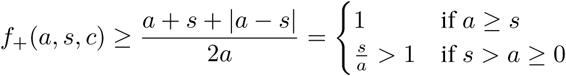

*Proof of 2 and 3.* When *a* < 0,

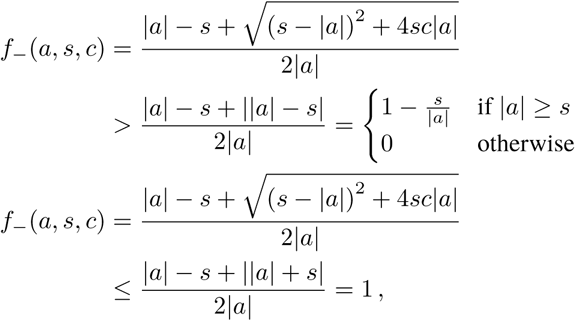

and the last inequality is strict when *c* < 1. When *a* > 0,

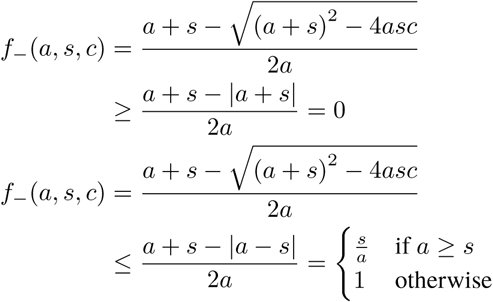

where we’ve used Eq. (E3). Note that that inequality is strict when *c* < 1. Then, 2 and 3 follow by combining and simplifying.

*Proof of 4.* We use L’Hôpital’s rule,

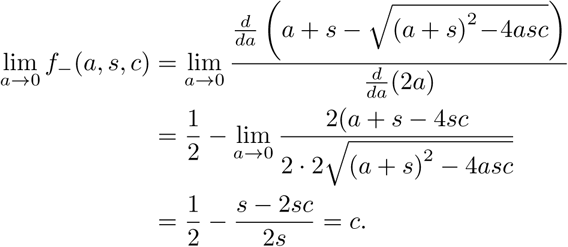

### Proof of 5.

If *c* = 1,

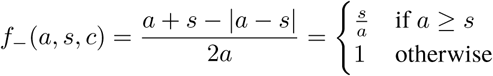

where we’ve used the proof of 2 and 3. Thus, when *c* = 1, 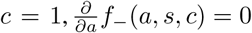 for *a* < *s* and 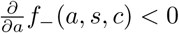 for *a* > *s*.

When *c* < 1, for notational convenience define

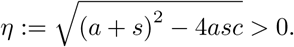

(The inequality comes from the fact that Eq. (E3) is strict when *c* < 1). Now, we consider the derivative,

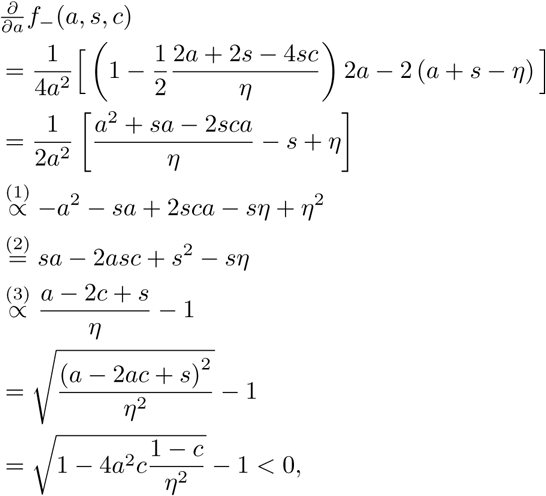

where in (1) we multiplied by the (positive) term 2*a*^2^*η*, in (2) we plugged in the definition of *η* and simplified, and in line (3) we divided by the (positive) term *ηs*. The inequality in the last line is strict since 0 < *c* < 1.

Using Lemma E.1, we now prove the following.

### Lemma E.2.

*Let c*(*x*) ∈ [0, 1] *indicate a set of values for all x ∈ 𝒜. Then, for any source distribution s_X_ with full support, there is a channel p_Y|X_ that satisfies*

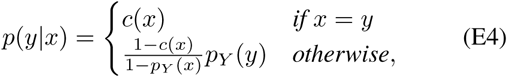

*where p_Y_ is the marginal p_Y_* (*y*) = ∑_*x*_*s*(*x*)*p*(*y*|*x*). *The channel is unique if c*(*x*) > 0 *for all x.*

### Proof.

We will show that there exists a marginal *p*_*Y*_ that satisfies the consistency conditions of Eq. (E4). The marginal is given by Eq. (E5), where *a* is a unique constant between 0 and 1 that makes ∑_*x*_*p*_*Y*_(*x*) = 1. Without loss of generality, we assume that there exists at least one *x* such that *c*(*x*) < 1 (otherwise, the result is trivially found by setting *p*_*YX*_ to the identity map).

We now proceed in several steps. First, we plug Eq. (E4) into *p*_*Y*_(*y*) =∑_*x*_*s*(*x*)*p*(*y*|*x*) to write

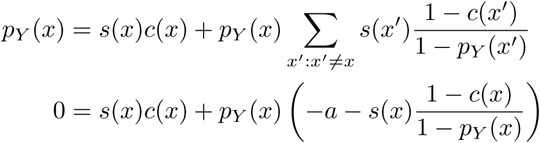

where we’ve defined the constant

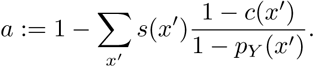

Multiplying both sides by 1 − *p*_*Y*_ (*x*) and simplifying gives

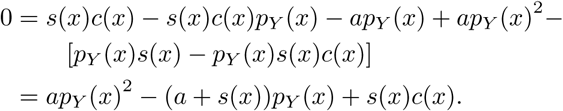

We solve this this quadratic equation for *p*_*Y*_ (*x*),

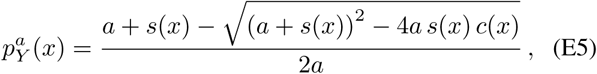

where we include the superscript *a* in 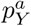 to make the dependence on *a* explicit, and we choose the negative solution of the quadratic equation. Our reason for choosing the negative solution are as follows:

- In case *c*(*x*) = 0, both branches may be valid and the solution is not necessarily unique, but for simplicity of analysis we choose the negative branch.
- In case *c*(*x*) > 0 and *a* ≤ 0, then by Lemma E.11, only the negative branch gives valid probability values.
- In case *a* > 0, then (by arguments that follow below) it must be the case that ∑_*x*_*c*(*x*) > 1, meaning that at least two *x*^′^, *x* ^″^ ∈ *𝒜* have *c*(*x*′) > 0 and *c*(*x*″) > 0 Then, since *s*_*X*_ has full support, both

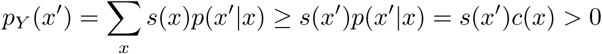

and (by a similar derivation) *p*_*Y*_ (*x*^*″*^) > 0, thus no *p*_*Y*_ (*x*) can be equal to 1. Thus, again by Lemma E.11, only the negative branch is valid, since it allows *p*_*Y*_ (*x*) < 1.

Note that these considerations imply that the solution is unique when *c*(*x*) > 0 for all *x*.

We now find *a*\*, the value of the value of *a* that satisfies 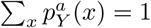. In other words, *a*\* is defined implicitly via:

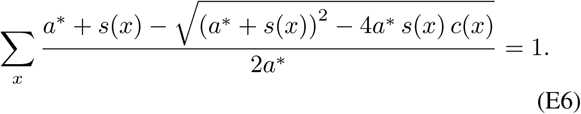

By Lemma E.12, Lemma E.13, and Lemma E.14,

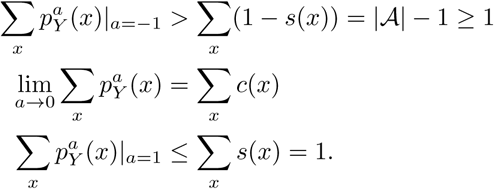

Moreover, 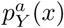 is continuous over *a* ∈ [−1, 0) and (0, 1]. This means that the *a*\* that satisfies Eq. (E6) must be:

1. within [−1, 0) if ∑ *c*(*x*) < 1,
2. equal to 0 if ∑_*x*_ *c*(*x*) = 1,
3. within (0, 1] if ∑_*x*_ *c*(*x*) > 1.

(The first and last claims follow from the Intermediate Value Theorem.) Finally, using Lemma E.15, we have

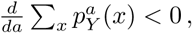

where we’ve used the assumption that at least one *x* has *c*(*x*) < 1. Thus, the *a*\* that satisfies Eq. (E6) must be unique.

In practice, the value *a*\* in the proof of Lemma E.2 can be found by a numerical root finding algorithm, or by trying values from − 1 to 1 in small intervals and selecting the first value that makes the LHS of Eq. (E6) less than or equal to 1. The marginal *p*_*Y*_ and channel *p*_*Y*_|_*X*_ can then be computed in closed form using Eqs. (E4) and (E5).

We now prove the following theorem.

### Theorem E.1.

*Let c*(*x*) ∈ [0, 1] *indicate a set of values for all x* ∈ *A, and s_X_ any source distribution with full support. If and only if* ∑_*x*_ *c*(*x*) ≥ 1, there is a channel p_*Y|X*_ *with diagonals p*_*y|x*_*(x) = c(x) and* 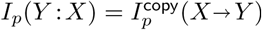

### Proof.

It is easy to verify that a given channel *p*_*Y|x*_ *with* given diagonals *c*(*x*) and 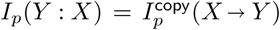 if and only if it has the form of the channel constructed in Lemma E.2, and all *c*(*x*) ≥ *p*_*Y*_ (*x*) for all *x*.

We now show that *c*(*x*) ≥ *p*_*Y*_ (*x*) for all *x* if ∑_*x*_ *c*(*x*) ≥ 1, and *c*(*x*) < *p*_*Y*_ (*x*) otherwise. As in the proof of Lemma E.2, we assume that we are not in the trivial case where all *c*(*x*) = 1.

In the proof of Lemma E.2, we found that the *a*\* that satisfies Eq. (E6) obeys *a*\* < 0 if ∑_*x*_*c*(*x*) < 1, and *a*\* ≥ 0 otherwise. At the same time, by Lemma E.15, 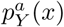 is strictly decreasing in *a* and, by Lemma E.15,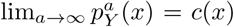. Thus, if ∑_*x*_*c*(*x*) < 1 it must be that *a*\* < 0 and 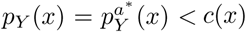.If ∑_*x*_*c(x)* ≤ 1 then a* ≥ 0 and 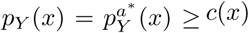.

We are now ready to prove Proposition 1.

### Proposition 1.

*For any source distribution s_X_ with full support and H*(*s*_*X*_) < *∞, there exist channels p for all levels of mutual information I*_*p*_(*Y* : *X*) ∈ [0, *H*(*s*_*X*_)] *such that* 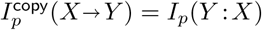

### Proof.

First, consider some source distribution *s*_*X*_ and mutual information level *I*_*p*_(*Y* : *X*) = *k*. We begin by showing that *p*_*YX*_, as defined in Lemma E.2 and Theorem E.1, is a continuous function of *s*_*X*_ and 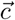 (in this proof, we use the notation 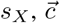 to indicate the vector of values *c*(*x*) for all *x*). Without loss of generality, we assume that 0 < *k* < *H*(*s*_*X*_), since it is trivial to find solutions for these edge cases (e.g., the channel *p*(*y x*) = *s*_*X*_ (*y*) for *k* = 0 and the identity channel *p*(*y*|*x*) = *δ*(*y, x*) for *k* = *H*(*s*_*X*_)).

Consider the channel *p*_*Y|X*_ from Lemma E.2. This channel is defined in terms of *s*_*X*_, 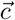, and *a*\*. In turn, *a*\* is implicitly defined in terms of *s*_*X*_ and 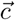 in Eq. (E6). We rewrite this implicit definition as implicit definition as

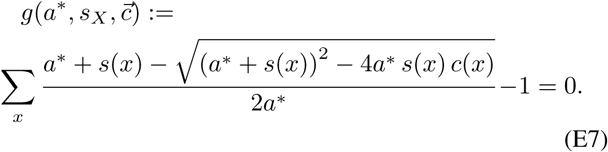

As shown in the proof of Lemma E.2, for a fixed *s*_*X*_ and 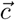 (assuming that Σ_*x*_ *c*(*x*) *>* 1 and there is at least one *x* with 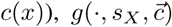 is strictly decreasing, and thus invertible. By the implicit function theorem, that means that the *a*\* that satisfies Eq. (E7) is a continuous function of *s* _*X*_ and 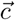 Thus, *p*_*Y*|*X*_ is a continuous function of *s*_*X*_ and 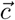.

Next, define a m anifold o f *c* (*x*) v alues p arameterized by *α* ∈ (0, 1) as

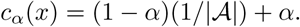

For a given source *s*_*X*_ and set of values *c*_*α*_(*x*) for a given *α*, let 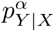 indicate the corresponding channel, as defined in Lemma E.2. Observe that for *α* = 0, 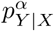 is completely noisy and thus 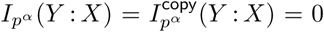. At the same time, for *α* = 1, the corresponding channel 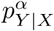 is the identity map, and it is easy to verify that 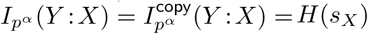.

Note that, for a fixed source distribution *s*_*X*_, mutual information is continuous in the conditional probabilities specified *p*_*Y*|*X*_. Clearly, *c* _*α*_ (*x*) is continuous in *α* and (as we’ve shown above) 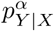 is continuous in *c*_*α*_ (*x*). Thus, the mutual in-formation 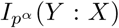 is continuous in *α* ∈ (0,1) (so that Σ_*x c* *α*_ (*x*) > 1), and must sweep all values in (0, *H*(s_*X*_)) (for *α* → 0 and *α* → 1 respectively). Simultaneously, we know by Theorem E.1 that 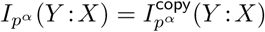

## Appendix F: The binary symmetric channel

The BSC is a channel over a two-state space (𝒜 = *{*0, 1*}*) parameterized by a “probability of error” ϵ ∈ [0, 1]. The BSC can be represented in matrix form as

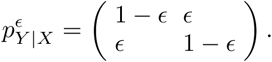

When ϵ = 0, the BSC is a noiseless channel which copies the source without error. In this extreme case, MI is large, and we expect it to consist entirely of copy information. On the other hand, when ϵ = 1, the BSC is a noiseless “inverted” channel, where messages are perfectly switched between the source and the destination. In this case, MI is again large, but we now expect it to consist entirely of transformation information. Finally, ϵ = 1*/*2 defines a completely noisy channel, for which mutual information (and thus copy and transformation information) must be 0.

For simplicity, we assume a uniform source distribution, *s*_*X*_ (0) = *s*_*X*_ (1) = 1*/*2, which by symmetry implies a marginal probability *p*_*Y*_ (0) = *p*_*Y*_ (1) = 1*/*2 at the destination for any *E*. For the BSC with this source distribution, Eq. (9) states that for both *x* = 0 and *x* = 1, 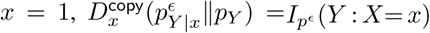 and 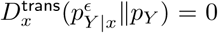 when ϵ *≤* 1*/*2, and 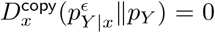 and 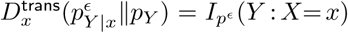 otherwise. Using the definition of the (total) copy and transformation components of total MI, Eqs. (13) and (14), it then follows that

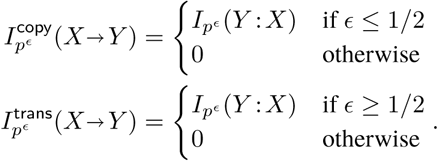

This confirms intuitions about the BSC discussed in the beginning of this section. The behavior of MI, *I*^copy^(*X* → *Y*) and *I*^trans^(*X → Y*) for the BSC with a uniform source distribution is shown visually in Fig. 2 of the main text.

